# SPA-STOCSY: An Automated Tool for Identification of Annotated and Non-Annotated Metabolites in High-Throughput NMR Spectra

**DOI:** 10.1101/2023.02.22.529564

**Authors:** Xu Han, Wanli Wang, Li-Hua Ma, Ismael Al-Ramahi, Juan Botas, Kevin MacKenzie, Genevera I. Allen, Damian W. Young, Zhandong Liu, Mirjana Maletic-Savatic

## Abstract

Nuclear Magnetic Resonance (NMR) spectroscopy is widely used to analyze metabolites in biological samples, but the analysis can be cumbersome and inaccurate. Here, we present a powerful automated tool, SPA-STOCSY (Spatial Clustering Algorithm - Statistical Total Correlation Spectroscopy), which overcomes the challenges by identifying metabolites in each sample with high accuracy. As a data-driven method, SPA-STOCSY estimates all parameters from the input dataset, first investigating the covariance pattern and then calculating the optimal threshold with which to cluster data points belonging to the same structural unit, i.e. metabolite. The generated clusters are then automatically linked to a compound library to identify candidates. To assess SPA-STOCSY’s efficiency and accuracy, we applied it to synthesized and real NMR data obtained from *Drosophila melanogaster* brains and human embryonic stem cells. In the synthesized spectra, SPA outperforms Statistical Recoupling of Variables, an existing method for clustering spectral peaks, by capturing a higher percentage of the signal regions and the close-to-zero noise regions. In the real spectra, SPA-STOCSY performs comparably to operator-based Chenomx analysis but avoids operator bias and performs the analyses in less than seven minutes of total computation time. Overall, SPA-STOCSY is a fast, accurate, and unbiased tool for untargeted analysis of metabolites in the NMR spectra. As such, it might accelerate the utilization of NMR for scientific discoveries, medical diagnostics, and patient-specific decision making.

## 1 Introduction

Many of the diseases that plague developed societies—obesity, diabetes, cardiovascular and neuro-degenerative diseases, cancer—are caused by or associated with faulty metabolic regulation. To measure composite metabolic changes, the medical community is increasingly relying on metabolomics (the study of all metabolites in a given sample) for quantitative phenotyping and biomarker discovery (Markley *et al*., 2017; Emwas *et al*., 2019; Psychogios *et al*., 2011; Wishart, 2019; Bujak *et al*., 2015; Newgard, 2017; Kennedy *et al*., 2018). However, metabolomics has still not been fully accepted as a critical tool for clinical assessment and mechanistic discoveries; the reasons mostly stem from cumbersome data analysis. Two main platforms used to acquire metabolomic data are mass spectrometry (MS) and NMR. MS is more sensitive than NMR; detecting more metabolites means gnon-destructive, and non-invasive; requires minimal sample preparation; and most importantly, it is quantitative (NMR signal intensity is proportional to sample concentration) at a dynamic range of 2×10^5^ and very reproducible (coefficient of variation, CV 1-2%) making it ideal for on-site diagnostics and biomarker discovery (Graaf, 2007). Because small changes in some metabolites (e.g. blood cholesterol, glucose) have serious health implications, identifying subtle-but-significant metabolic changes at the level of the whole metabolome can have a substantial clinical impact. Amassing relevant quantitative and highly reproducible data, such as those provided by the NMR, holds the promise to transform biology and medicine.

Identifying signals in NMR spectra may seem straightforward, but practical NMR spectral decomposition suffers from several challenges (Graaf, 2007). First, NMR data analysis is operator-based and time-intensive due to the complex nature of the data: the spectra contain thousands of resonances that may belong to hundreds of metabolites and although each metabolite has a unique spectral pattern, peaks of different metabolites may overlap. Second, real sample signal line shapes may differ from the idealized signals in the reference library, due to instrumental imperfections during acquisition. Third, a real sample may contain components not found in the library: a metabolite that is normally present at undetectable levels, or a breakdown product of an uncommon food or a rare drug (Graaf, 2007). Thus, NMR spectra may not correspond, precisely, to the weighted sums of ideal reference library components. Analyzing NMR spectra is also difficult due to high dimensionality: they have thousands of variables (data points) corresponding to thousands of dimensions, which can be computationally intense. Yet, NMR signal analysis methods should not attempt to simplify these computations by making assumptions about metabolites’ number and identity, as it is critical to extract the actual chemical, structural, or metabolic pathway information. For all these reasons, it is still challenging to use NMR for metabolomics discoveries.

To help surmount these challenges, several computational tools have been developed to perform standard data preprocessing and to obtain phased, baseline-corrected, chemical shift-referenced, and normalized spectra (van Beek, 2007; Ludwig and Günther, 2011; Günther *et al*., 2000; Worley and Powers, 2014; Pang *et al*., 2021). Existing computational and statistical tools for metabolomic profiling can then be applied to the processed spectra to extract the biologically useful information(Zheng *et al*., 2011; Hao *et al*., 2012; Lewis *et al*., 2009; Xia *et al*., 2009; Hollywood *et al*., 2006; Zhang *et al*., 2015). However, these methods, led by Statistical Total Correlation Spectroscopy (STOCSY) (Cloarec *et al*., 2005) and its variations (Robinette *et al*., 2013), have yet to accurately identify metabolites from the NMR spectra because of the high-dimensional problem in the covariance matrix estimation and the overlapping signals. In a typical NMR spectrum, the number of chemical shift intervals (ppm values) can range from 3000 to 8000 and the number of samples can range from several to hundreds. In addition, adjacent points on a spectrum often demonstrate high correlation, which makes it hard to estimate distal dependence. Further attempts have been made to first group NMR signals in a certain way and then utilize STOCSY: Cluster analysis statistical spectroscopy (CLASSY) employs a correlation matrix of peaks and an intersection matrix to determine local clusters and explore the intrametabolite connections (Robinette *et al*., 2009). Consequently, it identified the intermetabolite connections by hierarchical clustering of those local clusters. Statistical recoupling of variables (SRV) groups variables by scanning the covariance/correlation ratio landscape in consecutive variables. It then combines the grouped variables into super-clusters which represent variables with similar physical, chemical, and biological properties (Blaise *et al*., 2009). STOCSY is then applied to the super-clusters to help metabolite identification. Besides, there are also continuous efforts in the extension of STOCSY to accelerate the metabolites identification process. STOCSY-scaling method scales down the contribution of the metabolites with dominant intensities, such as glucose, which could explain most of the variation between the biological groups (Maher *et al*., 2012). It makes it possible to explore and identify metabolites that have the metabolic variation of diagnostic interest but are covered by those dominant metabolites. STORM is developed to advance biomarker discovery by applying STOCSY on selected subsets of spectra that contain specific spectroscopic signatures differentiating between different human populations (Posma *et al*., 2012). More recently, RED-STORM, the extension of STORM in a probabilistic framework, is applied for metabolite identification in 2D NMR data (Posma *et al*., 2017). Also, POD-CAST, which is applied after the STOCSY, aims at overcoming the peak overlap issues and provides a better peak list for database queries (Hoijemberg and Pelczer, 2018). Despite all these efforts, metabolite identification remains time-consuming and involves highly specialized knowledge and complex procedures on the part of the operator, while still failing to identify novel metabolites in the spectra.

Here, we present a novel, automated approach that identifies metabolites with spectral patterns in a known database (reference library) and provides insights into those that are not in the reference library. We took advantage of two existing algorithms: Spatial Clustering Algorithm (SPA) and STOCSY (Cloarec *et al*., 2005) (Fig. 1). SPA takes advantage of the strong correlations among data points from the same multiplets and identifies local spatial clusters of contiguous data points that are highly possible to arise from the same metabolite cluster. With the SPA-derived clusters as input, STOCSY is then used to highlight the correlation map and to arrange clusters into highly correlated groups containing signals from the same metabolite. Once combined, the analytical power of these two algorithms synergistically increases and compared to the existing methods, enables automatic, accurate, and fast identifications of metabolites in the spectra.

**Fig. 1.**
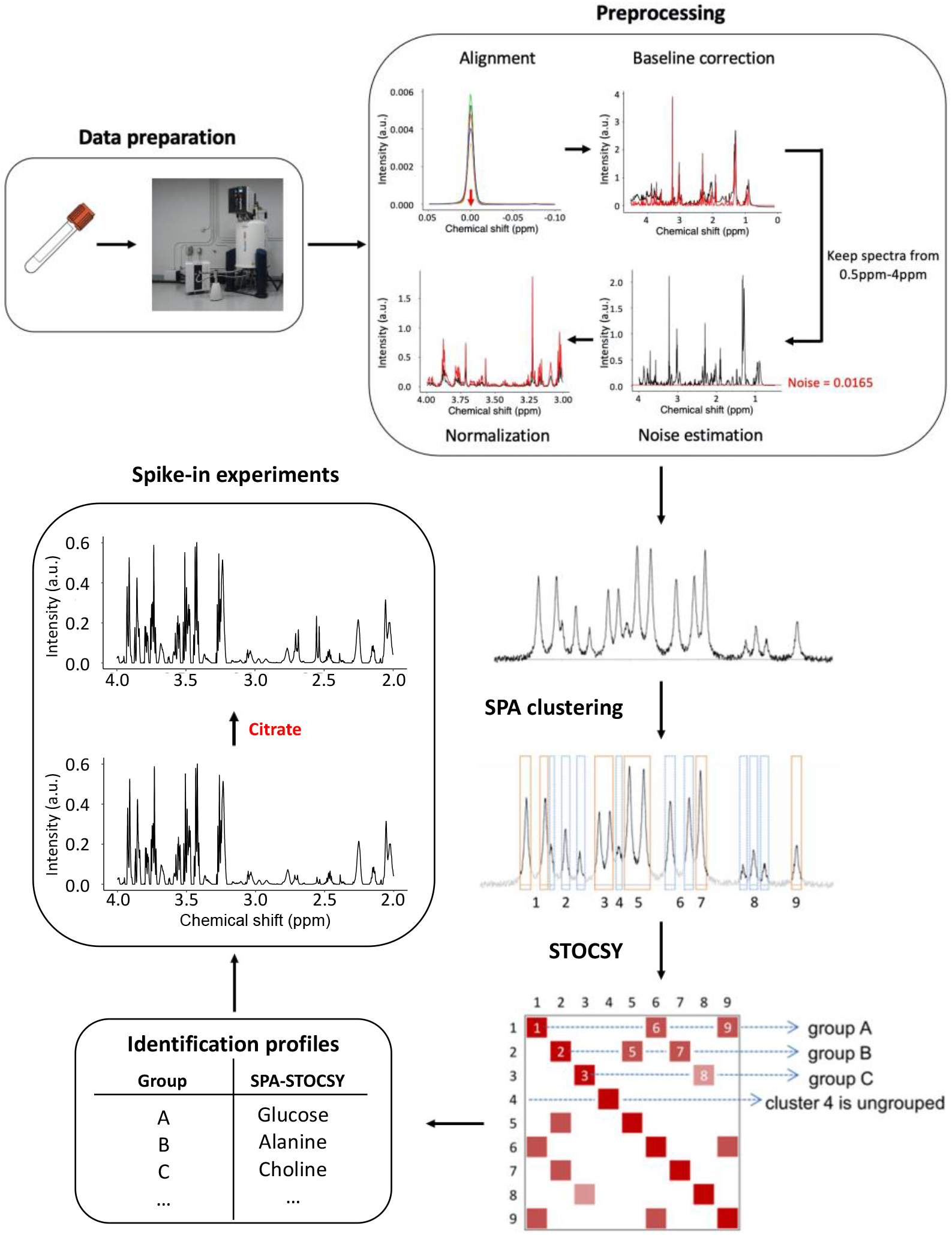
SPA-STOCSY flow chart. A set of spectra are experimentally acquired by NMR and the raw data are preprocessed. The mean spectrum of preprocessed spectra is shown before and after the SPA clustering. SPA automatically identifies clusters (outlined in orange or blue) that correlate strongly across multiple spectra (N>8). STOCSY of these clusters then automatically generates groups of clusters predicted to be from the same metabolite, i.e. the ^1^H chemical shifts of the clusters in each SPA-STOCSY group are predicted to be most likely from the same metabolite. Using the Chenomx, Inc. database as a reference library, SPA-STOCSY can generate an identification profile for each metabolite by summarizing the information from each cluster. Finally, to test the authenticity of identified metabolites, spike-in experiments can be performed. SPA=SPatial clustering Algorithm, STOCSY=Statistical Total Correlation Spectroscopy.

## 2 Methods

### 2.1 NMR simulation model

We generated spectral intensities as

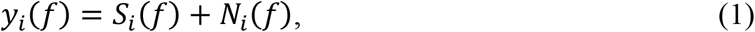

where *f* denotes the data points in terms of ppm, *f* = 1, …, p, and *i* denotes the samples, *i* = 1, …, *n*. A typical simulated spectrum with 50 metabolites is shown. (Supplementary Fig. 1a). We specified the noise term *N*(*f*) as an autoregressive process of order 1, i.e., AR(1), i^th^ high correlation (in simulations we used *ρ* = 0.9) (Supplementary Fig. 1b). This choice of the error term seemed reasonable based on the characteristics of the real data, and it has also been adopted in simulation models for other high-throughput spectral data, such as mass-spectrometry data (Cruz-Marcelo *et al*., 2008). Let *L* be the total number of metabolites present in a sample. *M*(*f*) is defined as reference spectra (scaled to have maximum 1) that the metabolite *l* resonates at the chemical shift, *f*, that is *M*_*l*_(*f*) ∈ [0, 1] (Supplementary Fig. 1c), and *Σ* ∈ *R*^*L*×*L*^ as a correlation matrix, denoting the correlation between the metabolites in the sample. The matrix *S*_(*n* × *p*)_ of population spectra, with elements *S*_*i*_(*f*). *S* is defined as

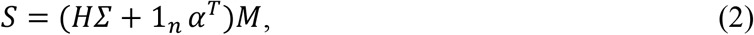

where *M* ∈ *R*^*L*×*p*^ has elements *M*_*l*_(*f*), *H* ∈ *R*^*n*×*L*^ has elements *H*_*i*_ ∈ ^*iid*^*N*(0, ∅^2^), denoting the variation of metabolite *l* in the *i*^*th*^ sample and *α* ∈ *R*^*L*×1^ such that *α*_*l*_ ∈ ^*iid*^*X*^2^(*γ*), denoting the mean concentration of each metabolite in the sample.

In simulations, we fixed *L*, the total number of metabolites in the samples, and *M*, the metabolite reference spectra. We then generated different datasets with different values of *n*, the sample size, *γ*, the degree of freedom for generating metabolite concentrations, and ∅, the variance of generating metabolite variation. These parameters, in turn, determined the signal-to-noise ratio (SNR), calculated as the variance of signals divided by the variance of noise: 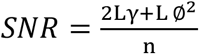. Parameter values were chosen to imitate experimental data from our lab, acquired on an 800MHz NMR spectrometer (Bruker, Inc). In particular, the mean value to generate metabolite concentrations and the variance for simulation were determined based on summary statistics from data that included both *in vivo* and *in vitro* experiments, acquired from several species such as humans, mice, and fruit flies. Means and variances of the intensities at the 25^th^, 50^th^, and 75^th^ percentiles were calculated from this experimental data. Simulation parameters were set to have summary statistics of the simulated data matching this range of values.

### 2.2 Simulation setup

We designed three sets of synthesized spectra, in which the total number of metabolites in the sample, *L*, was fixed at 10, 30, or 50 metabolites, respectively. The reference spectra were chosen from the BMRB database (Ulrich *et al*., 2008). The spectra were generated for the chemical shift range of 0.5 to 4.0ppm. Each reference spectrum was scaled to its maximum intensity so that each spectrum was within the range (0, 1). The scaled spectra were then used to construct the matrix *M* in the three simulated scenarios, according to the different values of *L*. We generated data according to models (1) and (2). In all simulations, we set *Σ* = *I*. We set the window size to *k* = 5, fixed *γ* = 60 and had either ∅ = 12 or 25. PACF was used to examine the autocorrelation in the dataset and determined *k* in a manner that at the *k*^*th*^ lag, the partial autocorrelation first falls into the 95 percent confidence interval. We performed analysis on two sample sizes (*n* = 50, 100). Each dataset (*n* = 50 or 100 samples per dataset) had a different combination of concentrations of the reference metabolites. However, the mean intensity of each spectrum was always the same.

### 2.3 Spatial clustering algorithm (SPA)

The first step of the SPA is to generate a correlation landscape since variables from the same functional unit tend to have stronger correlations with each other than variables from different functional units or the baseline (Fig. 2a, b). To achieve an accurate record of the relationships among individual variables, we designed a *moving average approach* to scan the correlations among consecutive variables. This moving average calculates *the correlation landscape* as the average pair-wise Pearson correlation between adjacent variables in a window of fixed size. Let us define

**Fig. 2.**
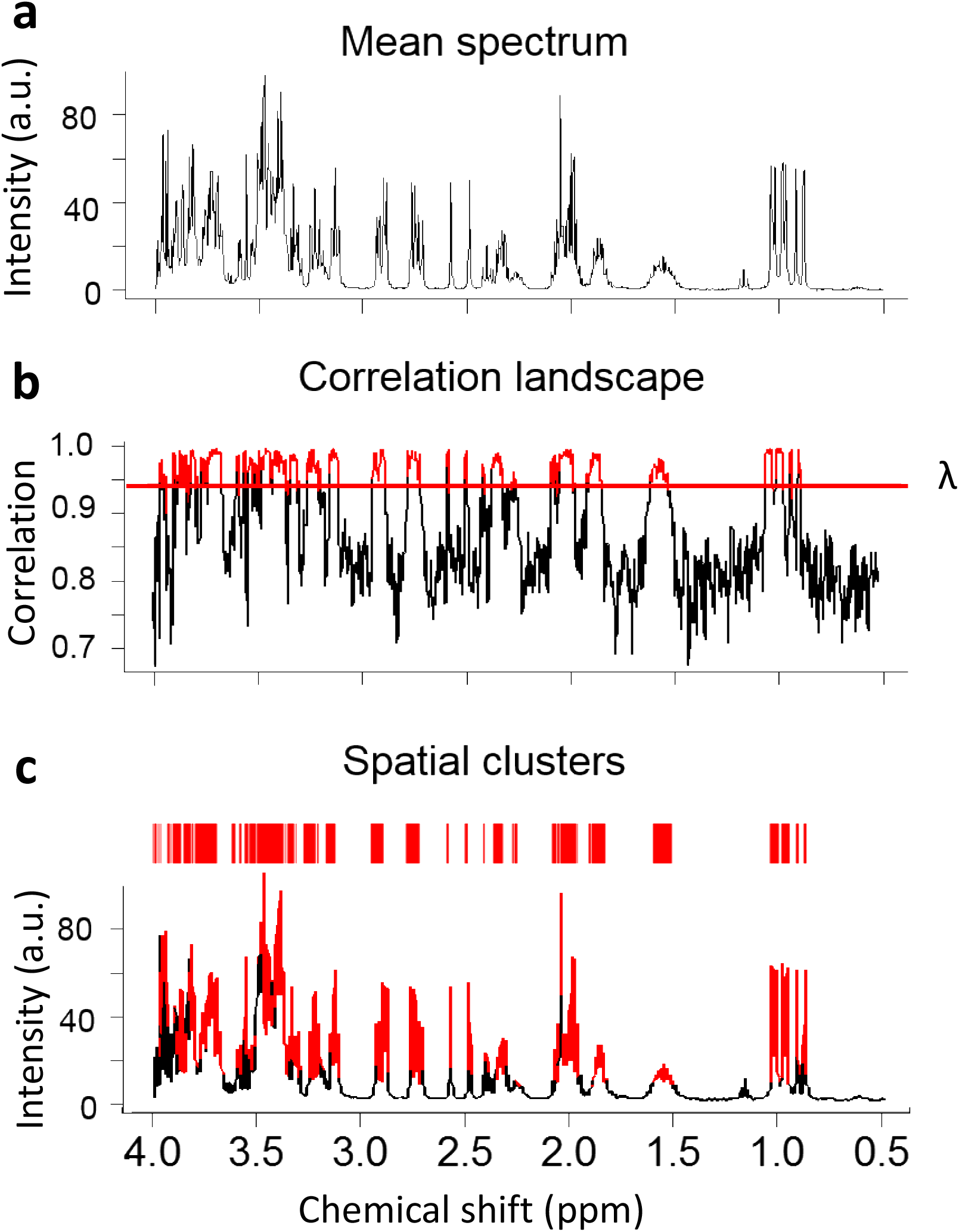
SPA Flowchart. **a**. The mean spectrum of the simulated NMR spectra of ten metabolites. **b**. Correlation landscape and the threshold λ. **c**. Identified SPA clusters putatively belong to the same structural units of a metabolite. Red indicates clustered regions.

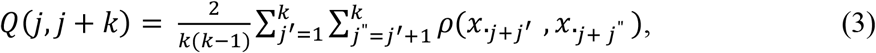

where *Q* is the correlation landscape, *j* is the index of the first variable in the window, and *k* is the window size. *ρ*(.) denotes the Pearson correlation coefficient. *X* denotes the dataset under study, whose columns represent the data points at each ppm value and rows represent sample.

Next, we improved the stability of the partial correlation landscape by kernel smoothing. Based on our simulated data, the Epanechnikov kernel,

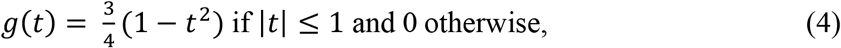

and the tri-cube kernel,

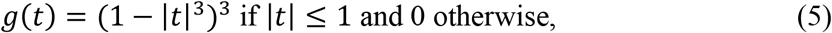

are best suited for spectral data. To identify SPA clusters in a spectrum (Fig. 2c), we need to set a threshold on the correlation landscape λ. Although this threshold can be empirically determined, we set it by optimizing the correlation thresholds that maximizing the stability for ppm memberships between two equally split datasets out of *X* (the whole dataset under study). We designed an automated procedure based on the prediction strength (Tibshirani and Walther, 2005). It is calculated as the median proportion of correctly allocated variables across all spatial clusters in a sampled set (Tibshirani and Walther, 2005). This procedure starts by first randomly splitting the data into two sets of equal sample size, set-1, and set-2. Our goal is to find the threshold *λ*, at which the spatial clusters become most stable between set-1 and set-2. At the same time, the grouped variables should be as informative as possible to represent the chemical structural information of the present metabolites. To accomplish this goal, a *p* × *p* co-membership matrix *D*, representing how well the memberships in set-1 can predict the memberships in set-2, is constructed. If variable *i* in the set-1 and variable *i*_0_ in the set-2 are in the same cluster, then *D*(*i, i*_0_) equals 1, otherwise 0. Let *c* be the total number of clusters, *m*_1_, …, *m*_*c*_ denote the number of variables in each spatial cluster and *C*_1_, …, *C*_*c*_ are the spatial cluster memberships. For a given threshold *λ*, the prediction strength *Pd* is defined as the median proportion of correctly allocated variables, across all spatial clusters in the set-1.

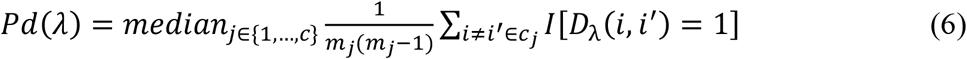

This process is repeated for several random folds to determine the mean and standard error of the prediction strength. The optimal threshold is then chosen as the lowest value whose mean prediction strength is within one standard error of the maximum mean prediction strength over all possible thresholds.

### 2.4 Statistical Recoupling of Variables (SRV)

Let *X*_*n*×*p*_ denote the data matrix, with rows representing samples and columns representing spectral variables. Let us define

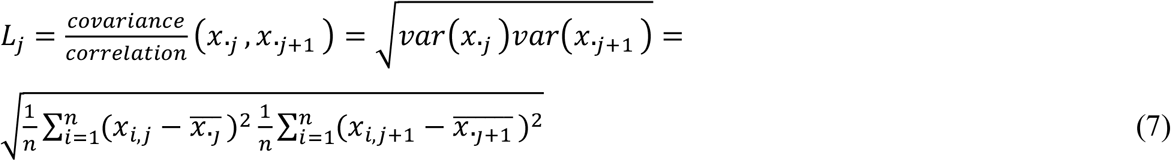

for *j* = 1, …, *p*. The boundaries of spectral clusters are identified by the local minima of the landscape function. The minimum number of variables in one cluster is determined by the resolution of the NMR spectra. Blaise et al. suggest discarding clusters that have less than 10 variables (Blaise *et al*., 2009). Representative intensities of spectral clusters are calculated as the mean of NMR signals assigned to each cluster. Super-clusters are then formed by aggregating clusters according to their correlation with neighboring ones. The authors suggest aggregating clusters with the correlation greater than 0.9 and limiting the association to a maximum of three clusters to keep efficiency in areas of highly correlated signals.

### 2.5 SPA input for STOCSY

To couple these spatial clusters with STOCSY, we first calculated each cluster’s intensity as a mean value of the local-maximum peaks within the cluster and generated a new data matrix. We input this data matrix into STOCSY. Using 0.8 as the threshold for SPA-STOCSY, we identified the highly correlated clusters. We then found these clusters in the spectrum and determined the corresponding resonances. For both known and unknown metabolites, the identification can be realized using these distinguished signals and their resonances either with a reference library or by fragment-matching using the identified splitting pattern.

### 2.6 NMR sampling

The *Drosophila* head tissues (Control, Elav-GAL4/w1118; +; +) and human embryonic stem cells (hESCs) (1 million cells) were each dissolved in phosphate-buffered saline, pH 7.25, containing 10% D_2_O as a field frequency lock. One-dimensional ^1^H-NMR spectra were collected at 23^0^C using an 800 MHz NMR spectrometer (Bruker Inc.). DSS (4,4-dimethyl-4-silapentane-1-sulfonic acid) was used as the internal standard for head tissues, while for hESCs we used 0.05mM TSP (Trimethylsilylpropanoic acid) as the chemical shift internal standard.

For *Drosophila* head tissues, for each free induction decay (FID), 128 transients were recorded at a repetition rate of 2 sec per transient. To minimize the large water peak, the water signal was pre-saturated with a low power radiofrequency (RF) pulse. For hESCs, to minimize the water peak, the ZGESGP pulse sequence from the Bruker library was selected. For each FID, the size of FID is equal to 32K, the number of dummy scans is 4, and the spectral width was 12ppm. 128 transients were recorded at a repetition rate of 1.7 sec per transient.

Finally, spectra from both datasets were preprocessed in Topspin including Fourier transform to the frequency domain, Gaussian filter, centering of the water peak to 4.7ppm, binning, phasing, baseline correction, peak alignment to the internal standard, and the normalization to the integral of the spectrum were performed. To remove the dilution factor in samples, we applied the probabilistic quotient normalization on the normalized dataset (Dieterle *et al*., 2006). Similar to others, we observed that the variables at baseline regions in the preprocessed data had notably high correlations. We estimated the baseline level by calculating five standard deviations from a noise region of the baseline and set the points with intensities less than the baseline level to zero, as suggested by Torgrip et al (Torgrip *et al*., 2008). Here, we considered the noise region as 0.08-0.58ppm.

## 3 Results

### 3.1 SPA allows the identification of strongly correlated functional units

In the NMR spectrum, each metabolite has unique signatures depending on its chemical and physical structure. As noted, the signature is represented by one or more structural units, which can be singlets, doublets, or multiplets in the spectra. The SPA identifies these structural units even where many of the signals overlap (Fig. 2) because unlike existing clustering methods, which focus on individual peaks or the variation of peaks, the SPA aims to group data points that belong to the same structural unit. It does so by finding contiguous groups of data points that exhibit stronger correlations since data points from the same multiplets correlate more strongly with each other than with those from other multiplets.

First, the SPA seeks to determine the relationships among individual data points. After preprocessing, a typical NMR spectrum has more than 3,000 data points which can generate about 5,000,000 pairs of correlations, each of which contain complex patterns. To calculate the correlation landscape, the SPA first determines the window size, which is a type of bucketing. Bucketing is commonly used to control the peak shifts in NMR data, but the drawback is a loss of resolution (Forshed *et al*., 2003). High-resolution bucketing is therefore needed but decreasing the size of the bucket can cause the loss of spectral dependence (Blaise *et al*., 2009). Accordingly, the SPA uses the partial autocorrelation function (PACF) approach to determine the window size, thus balancing resolution quality and spectral dependence.

To illustrate, assume a data point at location *n* and another data point at location *n-k*. The partial autocorrelation of them is the correlation of the two data points conditionally on the data points between them. This approach estimates a window size *k* that data points within this window will have significantly stronger dependencies.

To test the performance of the SPA, we first synthesized a spectrum using 10 random metabolites with complex spectra (Supplementary Fig. 1). After a series of calculations and filtering, the SPA-produced spatial clusters contained data points and peaks with the strongest correlation and without noise or baseline data points (Fig. 2b, c). Therefore, these clusters were more likely to be functional units, representing the structural information of the underlying molecules.

### 3.2 SPA consistently outperforms SRV in discriminating signals from noise

We then compared the SPA to SRV (Blaise *et al*., 2009), another clustering method that calculates a landscape function as the ratio of covariance and correlation between neighboring spectral data points. We were prompted to compare their performance because the two methods grouped data points of the same simulation dataset differently (Fig. 3a).

**Fig. 3.**
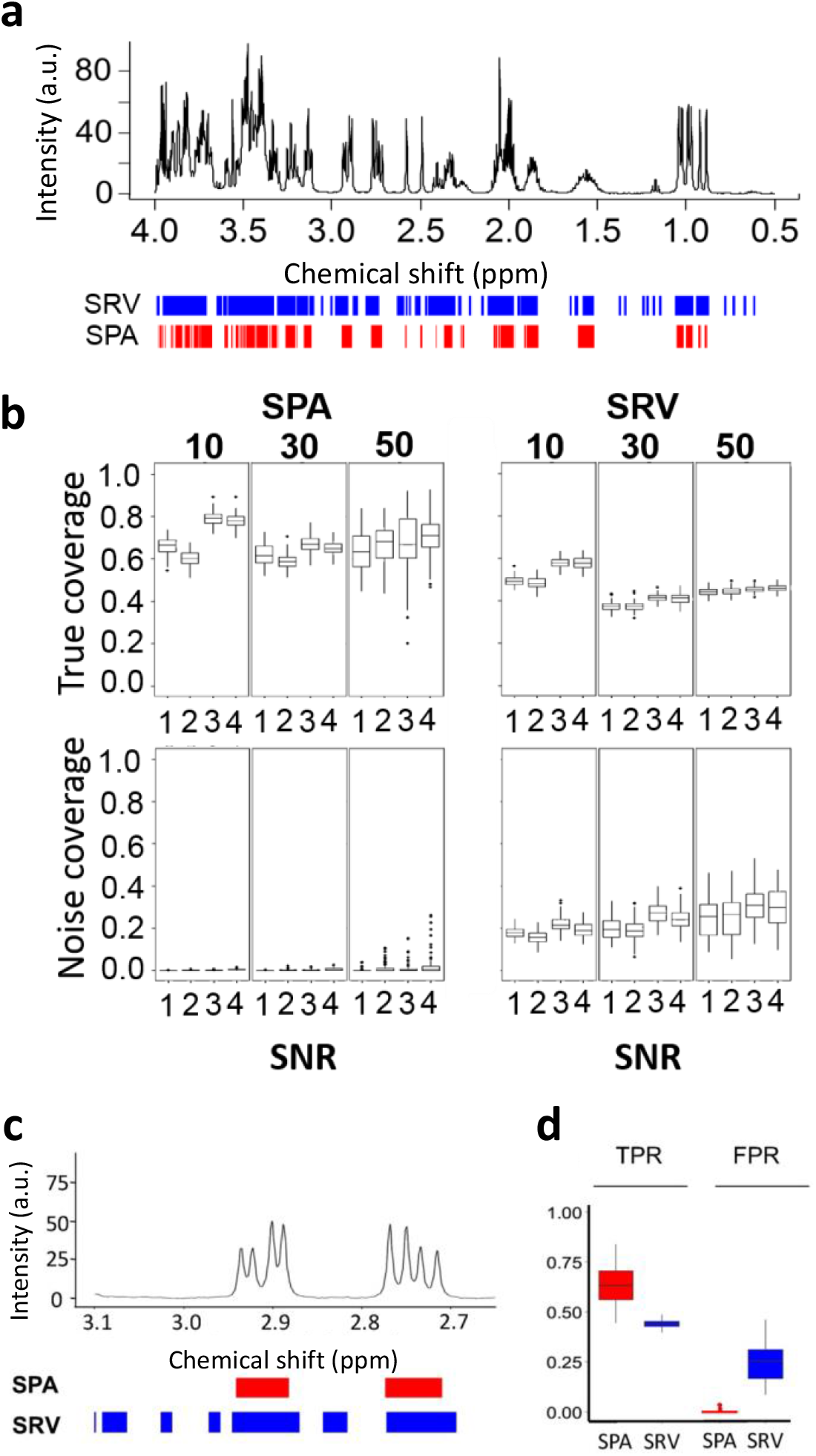
Comparison of the SPA and SRV clustering performance. **a**. SPA (red) and SRV (blue) clustering on a simulated spectrum containing 10 metabolites. **b**. The performance of each method was measured. *True coverage*: Proportion of detected true metabolite regions, measured as the percentage of the metabolite regions detected over the true metabolite resonance regions. *Noise coverage*: Proportion of detected regions in which no metabolite resonates. This is measured as the number of detected variables located in noise regions, divided by the total number of noisy variables. Boxplots show the true coverage and noise coverage from simulated datasets using SRV and SPA of all the simulation scenarios (Supplementary Table 1). In each scenario, 100 simulations were conducted. **c**. A synthesized spectrum containing 50 metabolites was analyzed by both methods. SPA discriminates between signals and noise better than SRV, as illustrated on a zoomed region of the spectrum. Boxes indicate assigned clusters (red: SPA; blue: SRV). **d**. Boxplots show the true positive rate (TPR) and false positive rate (FPR) for SPA and SRV. SRV=Statistical Recoupling of Variables.

We designed three sets of simulated spectra, in which the total number of metabolites in each sample was fixed at 10, 30, or 50, respectively. (The more metabolites in a sample, the greater coverage of overlapping regions where metabolites resonate.) For each simulated set, we generated four scenarios by varying metabolites’ concentrations, sample size, and sample variations (Supplementary Table 1). With these different simulation parameters, each scenario had a different SNR (Supplementary Table 1). When analyzing the spectra, an ideal method would demonstrate high *true coverage* (detection of true metabolite regions) and low *noise coverage* (detection of regions in which there is no metabolite). For each scenario, we calculated the SPA’s and SRV’s true coverage and noise coverage (Fig. 3b) and ran the analysis 100 times for each sample (for example, for a dataset with 50 samples, the total number of tested samples was 50 × 100). Increasing the total number of metabolites in the sample resulted in increased variability of both true and noise coverage for the SPA, but only noise coverage for SRV. As expected, the true coverage increased with increased SNR for both methods, while the sample size did not affect the results (Fig. 3b). Comparisons of the difference between the SPA and SRV measurements were all significant according to the student’s t-test (Fig. 3c, d). The SPA’s advantage over SRV is demonstrated by the clear difference in their respective true coverage and noise coverage ratios (Fig. 3). This finding is of critical importance because it indicates SPA reliably discriminates signals from noise.

### 3.3 Coupling SPA and STOCSY achieves accurate identification of both separated and overlapping metabolites on the simulated data

While the SPA can identify structural units in the spectra, it cannot by itself identify their corresponding metabolites. We, therefore, coupled the SPA with STOCSY, a method that uses statistical correlations between spectral data points to identify signals originating from the same molecule (Cloarec *et al*., 2005). STOCSY has been used previously to explore the intra- and intermetabolite correlations within raw NMR data, but high dimensionality and overlapping signals can make STOCSY readout and interpretation cumbersome. In SPA-STOCSY, however, we input the spatial clusters selected by SPA rather than raw NMR data, and the spectrum is therefore interpreted as a series of spatial clusters rather than data points. SPA-STOCSY then provides correlations between these spatial clusters, allowing the identification of corresponding metabolites.

We tested the SPA-STOCSY algorithm using simulated spectra with a set of 10 metabolites (spectrum shown in Fig. 2a). We first calculated each cluster’s intensity as a mean value of the peaks within the cluster and generated a new data matrix to be loaded into STOCSY. Using 0.8 as the threshold for SPA-STOCSY, we then identified the highly correlated clusters (Fig. 4a). Each set of highly correlated SPA clusters, when projected to the mean spectrum, displayed the reconstructed signature of the metabolites that might exist in the sample. For example, valine resonated in three clusters, which were accurately identified by SPA-STOCSY (Fig. 4b). To achieve this result, the SPA must effectively capture the true metabolite regions (Fig. 3d), and STOCSY must accurately pick up these clusters (Fig. 4a). When we input spectrum resonances that are highly correlated with clusters in the Biological Magnetic Resonance Data Bank (BMRB) database (Ulrich *et al*., 2008), SPA-STOCSY identified 9 out of 10 metabolites present in the simulated dataset (Fig. 4c). Thus, we conclude that the spatial clusters are close representations of the resonances from metabolites that exist in each sample. Moreover, STOCSY can correctly extract clusters that belong to the same molecule.

**Fig. 4.**
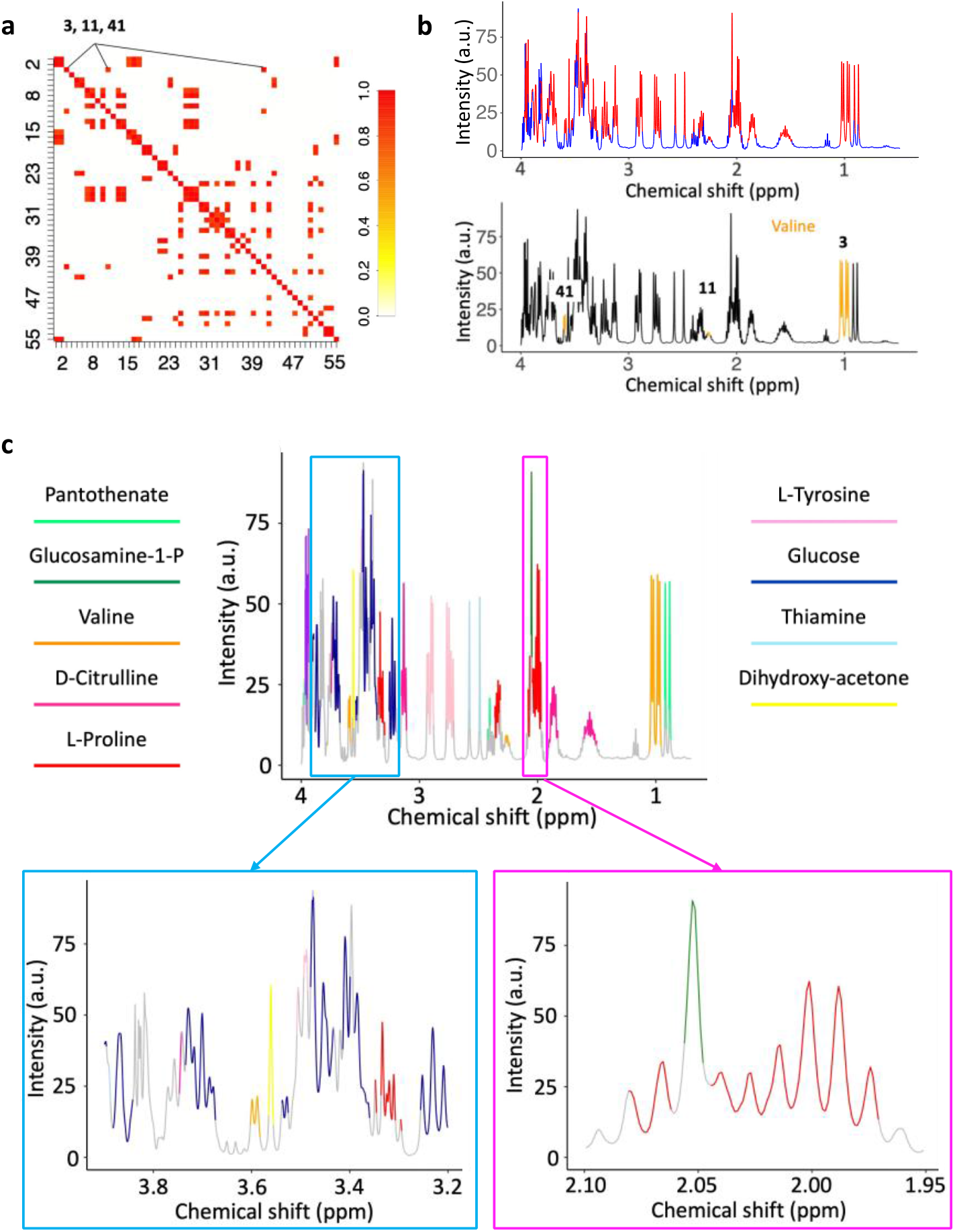
SPA coupled with STOCSY identifies connected fragments of molecules in the NMR spectra and thus, metabolites in the samples. **a**. STOCSY is performed on spatial clusters obtained from the simulation dataset with 10 metabolites (Fig. 2a). 55 spatial clusters are identified with a threshold of detection 0.8. **b**. SPA clusters spectra into spectral regions (red) and gaps (blue). The example is given for clusters 3, 11, and 41, identified by SPA-STOCSY as highly correlated and therefore likely to belong to the same metabolite. Indeed, they belong to valine. **c**. 9 out of 10 metabolites in the simulated dataset are correctly identified by SPA-STOCSY in a single pass. **d**. Zoomed-in regions from (**c**). *Left*: SPA-STOCSY discriminates highly overlapping regions between 3.2 to 4.0 ppm, where the method identifies glucose, dihydroxy-acetone, L-tyrosine, and valine peaks. *Right*: SPA-STOCSY discriminates highly overlapping regions between 1.95 to 2.10 ppm, where the method identifies L-proline and glucosamine-1-phosphate (a singlet embedded in an unrelated multiplet).

Upon inspection of the selected metabolites, we recognized that some reconstructed signal regions were incomplete, such as the glucose signal regions at 3.2 to 4ppm (Fig. 4d). This omission is caused by the overlapping signals from different molecules, which can reduce the existing correlations (Cloarec *et al*., 2005). However, as SPA-STOCSY identified the majority of glucose structural units, we were still able to identify this metabolite (Fig. 4d).

Peak overlapping is a universal challenge in NMR-based metabolomic profiling, and many algorithms lose accuracy when overlapping spectral peaks are present in a sample. But SPA-STOCSY substantially aids the identification of molecules because of its unique ability to identify overlapping peaks. In the SPA algorithm, the correlation landscape is sensitive to even minor changes in pair-wise correlations of the consecutive *k* data points, in which *k* is the window size. The fluctuation of the correlation landscape thus represents both the connectivity strength of high-correlation resonances that may originate from the same molecule, as well as the separation capability of resonances from different molecules or overlapping peaks (Fig. 2b). Spatial clusters tend to split where the overlapping occurs because it reduces the original strong correlations among data points originating from the same metabolite multiplet. When we apply STOCSY, the split clusters describing the same functional unit can be recombined because they are highly correlated, thus reconstructing metabolite regions. We demonstrate this property in two examples: L-proline and glucosamine-1-phosphate N-acetyltransferase, which overlap at the 2.0ppm region (Fig. 4e), and the glucose region between 3.2 to 4ppm, where the method identified glucose, dihydroxy-acetone, L-tyrosine, and valine peaks (Fig. 4d). These findings indicate that the overlapping problem may be overcome by correctly identifying and splitting the structural units in the overlapping regions. When correctly identified, structural units are then coupled in the STOCSY algorithm and the true metabolite regions are reconstructed.

### 3.4 SPA-STOCSY provides automatic identification of metabolites and outperforms current methods

The SPA-STOCSY algorithm has several automated features that quickly and accurately identify candidate metabolites and potentially avoid human interference in processing NMR data. These automated features include (1) determining window size for the calculation of correlation landscape; (2) deciding the threshold value for grouping data points that are likely from the same spatial cluster, according to the correlation landscape; and (3) identifying metabolites by matching highly correlated SPA clusters with the reference library. The first two parameters are determined automatically through exploration and investigation of the dataset under study. To connect the SPA-STOCSY data to the metabolite reference library, we designed an algorithm that provides a readout of all candidate molecules by matching spatial clusters resonances with the library’s reference resonances. Specifically, we identified each spatial cluster’s peaks using the local minimum and maximum for peak detection (Gong Lixin,Constantine William, 2012). Highly correlated spatial clusters can be transformed into a series of highly correlated peaks, which the algorithm then inputs into the reference library to find corresponding metabolites.

With these features, SPA-STOCSY outperforms current automated identification and visualization methods. For example, MetaboID requires user selection of candidate metabolites, which introduces operator bias (Mackinnon *et al*., 2013). MetaboHunter enables automatic identification of metabolites (Tulpan *et al*., 2011), but it matches individual peaks to the reference library rather than a group of peaks that originate from the same structural unit, resulting in a high false detection rate. MetaboHunter’s mutual exclusion of peaks also neglects overlapping signals, increasing the need to visually inspect the spectra for the metabolite readouts. BAYESIL segments spectra into small blocks and uses the probabilistic graphical model to estimate the metabolite concentrations and chemical shifts. But accurate quantification from BAYESIL is more applicable to specific biofluids with the sub-libraries curated from serum, plasma, and cerebrospinal fluid (CSF) (Ravanbakhsh *et al*., 2015). FOCUS is an integrated tool that provides a data analysis workflow to analyze 1D NMR-based metabolomics. Its metabolite identification concentrates on peak picking and matching, with the peaks extracted in pure compounds’ reference spectra (Alonso *et al*., 2014). Although FOCUS’s matching score mechanism considers peak correlations, with peak overlapping, its score is undermined due to a high proportion of zero-intensity peaks. ASICS is a linear-based tool whose accuracy depends heavily on a dedicated shifting algorithm that contains two steps: global shift and local peak distortions (Lefort *et al*., 2019; Tardivel *et al*., 2017). In the real application, the maximum allowed global shift needs to be well-tuned to reach the best alignment. This time-consuming adjustment could result from the different sensitivities among metabolites and sample heterogeneity. However, it makes it hard to set a standard among them, and it is inconvenient to tune them separately. Thus, while using residuals to find out the best distortions for peaks in metabolites, ASICS is likely to align reference spectra to the wrong clusters in heavily overlapping regions.

A biological sample is usually complex, and its total number and composition of metabolites are unknown. Moreover, the composition of metabolites varies in different samples and even scans acquired from replicated samples can be slightly different. One metabolite may present in most of the samples, but not in others. Despite this complexity, the connectivity of the underlying unknown signals from the same metabolite persists and can be captured. SPA-STOCSY uses covariance patterns from a dataset to recover such connectivity, thereby identifying a given sample’s missing signals that are present in many other samples in the same dataset.

### 3.5 Identification of metabolites in real data

We also tested SPA-STOCSY’s performance on real data, first with a set of experimental NMR data from the head homogenates of *Drosophila melanogaster* (12-day-old, female, 40 heads per sample, N=10) (Fig. 5a-c). Using PACF, we determined the optimal window size for scanning the correlation landscape as 7, and we chose a tri-cube kernel smoother to calculate the correlation landscape. Following the SPA, we identified 50 clusters that exhibited strong correlations (Fig. 5a) and observed that high correlation landscape values are always localized to signal regions. In addition, some regions had lower correlation landscape values and spatial clusters were split there. The sharp decline of correlation landscape values indicated overlapping, and the splitting behavior was more common in regions between 3 to 4ppm, abundant in overlapping resonances from different metabolites. The SPA can detect minute changes in correlations among variables resulting from overlapping peaks, leading to the accurate identification of those metabolites. We then analyzed the data with STOCSY. Here, the most important step was to determine the threshold of detection, i.e., the optimal correlation to provide the maximum number of meaningful variables. In real data, the optimal correlation threshold needs to be determined by the characteristics of the spatial clustering method itself. As the SPA groups variables that exhibit strong correlations, however, there is a chance that the correlations of highly concentrated metabolites at specific regions may overwhelm the metabolites present in low concentrations. Therefore, to ensure that SPA-STOCSY captured all correlated clusters, we took advantage of the external chemical reference, DSS. DSS resonates at several frequencies along the spectrum (Ulrich *et al*., 2008) and these J-coupled resonances have high correlations in SPA-STOCSY. Thus, the correlation threshold value when all DSS resonances were captured by STOCSY was chosen as the optimal value. Using this criterion, the optimal correlation threshold for the *Drosophila* head NMR dataset was 0.8.

**Fig. 5.**
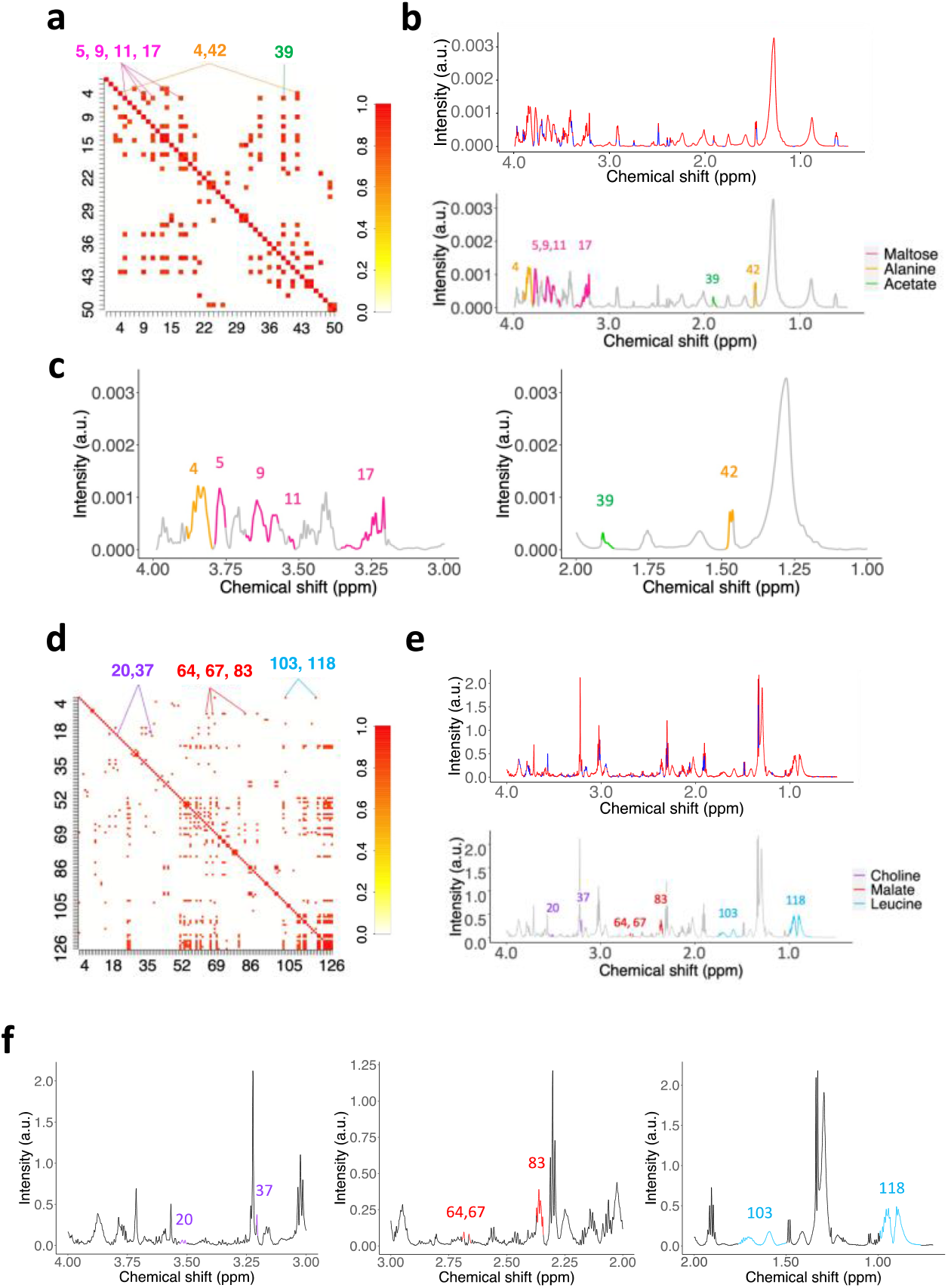
SPA-STOCSY identifies metabolites in the *Drosophila melanogaster* tissue and human cultured cells. **a**. SPA-STOCSY identifies 50 highly correlated clusters at a detection threshold of 0.8 in the *Drosophila* data. **b**. SPA clusters the mean spectrum (N=10) into structural units. The spectrum is color-coded by the correlation landscape values (red: strong correlations; blue: weak correlations). Clusters with high correlations are deconstructed into original resonance frequencies and the identities of the corresponding metabolites are obtained. Maltose, alanine, and acetate are given as examples. **c**. Amplified version for the visualization of highly correlated clusters and the corresponding metabolites from (**b**). **d**. SPA-STOCSY identifies 126 highly correlated clusters in human embryonic stem cell (hESC) datasets. **e**. SPA clusters the mean spectrum (N =22) into structural units. The spectrum is color-coded by the correlation landscape values (red: strong correlations; blue: weak correlations). Clusters with high correlations are deconstructed into original resonance frequencies and the identities of the corresponding metabolites are obtained. Choline, malate, and leucine are given as examples. **f**. Amplified version for the visualization of highly correlated clusters and the corresponding metabolites from (**e**). SPA-STOCSY accurately identifies molecules with complex signatures regardless of the overlapping and/or splitting regions.

Next, using a similar analysis, we applied SPA-STOCSY to 22 NMR spectra from hESCs. Using PACF, the window size was set at 11. The SPA generated 126 clusters in total. DSS resonances were barely captured due to the distortions and low concentrations, so to determine the optimal correlation threshold for this set of spectra, we took the two clusters from choline as the reference and set this threshold at 0.8 as well.

In the *Drosophila* head data, we set the automatic detection ratio at 0.55 and identified 132 metabolites using the Chenomx library as reference (Supplementary Table 2). Some examples of the correctly identified metabolites are presented in Figures 5a-c. As shown, the complexity of the resonances clustered by SPA-STOCSY ranged from 1 (acetate) to > 5 (maltose), including the regions of highly overlapping signals, such as between 3.00-4.00 ppm. We also performed an analysis of the same samples using commercially available software, Chenomx (Chenomx Inc., Edmonton, Canada). Chenomx analysis identified 81 metabolites (three of which were not in our library). To discern the differences, we visually inspected the identified metabolites, focusing on those that resonated from 0.00-4.00ppm. SPA-STOCSY and Chenomx identified 55 common metabolites, while SPA-STOCSY identified more metabolites missed by Chenomx. However, it is possible that SPA-STOCSY could capture false-positive metabolites because its automatic identification is based on the peak detection and cluster positions for metabolites in the reference library. Metabolites with only one cluster are likely to be claimed as detected. Thus, metabolites with multiplets captured by SPA-STOCSY offer stronger confidence (Supplementary Table 2).

For the hESCs, when the detection ratio was also set at 0.55, SPA-STOCSY identified 126 clusters and 81 metabolites. Three representative metabolites are shown in Figures 5d-f. Clearly, there is an increased level of NMR spectra complexity in the hESC samples as compared to the *Drosophila* samples. Nonetheless, SPA-STOCSY still captured highly correlated clusters from the same metabolite in diversified regions, with clusters ranging from singlets to multiplets. Chenomx profiling of the same hESCs data by an NMR expert identified only 24 metabolites. Compared to manual profiling, SPA-STOCSY, therefore, is more efficient and offers more testable candidates. For the 24 metabolites identified by Chenomx profiling, SPA-STOCSY detected at least one cluster for each of them if their clusters were included in the reference library within the selected region (0ppm to 4ppm) (Supplementary Table 3). For most of them, SPA-STOCSY achieved a detection ratio of at least 0.5. We thus believe that SPA-STOCSY is the first method to enable automated analysis of spectra with unknown number and type of metabolites with precision and speed (seven minutes).

## 4 Discussion

In this study, we have introduced a new algorithm, SPA-STOCSY, as an untargeted metabolome profiling tool. The SPA part of the algorithm greatly reduces the high dimensionality of the NMR dataset by transforming the data points into spatial clusters that represent functional units. First, SPA groups adjacent variables that potentially originate from the same functional unit and blindly identifies metabolites. In addition, it groups variables according to their average correlations, which can capture small changes in pair-wise correlations among variables within a determined window size, *k*, along the spectrum. Spatial clusters split where a sudden decline in average correlations occurs, as in the case of overlapping resonances. Therefore, the SPA aids the identification of metabolite regions by detecting overlapping components.

When analyzed by a human expert, the noise and overlapping peaks are the major factors that raise bias in metabolite identification. With SPA, we show its capability in making full use of the spectral information to limit the influence from a subjective bias. SPA outperforms SRV in successfully separating signal regions from noise regions. This clear signal-to-noise separation provides a good start for a human expert analysis and decrease bias. Also, later for the development of NMR quantification computational methods, the quality of the spectra is expected to be higher. SPA, which can deliver a noise-free spectrum within 2 minutes, can ease the preprocessing of real-world spectra without losing marginal spectral information.

With the wide applications of SPA, it has boosted performance when it is coupled with STOCSY. The significant feature of the SPA enables accurate metabolite detection, as shown in both our simulation studies and real data. For metabolites identification, a crucial task is to separate the overlapping peaks. With SPA itself, it can only separate the signals from noise, but not metabolite from metabolite. And for human experts, these separation boundaries are mostly decided based on their experience, raising the problem of subjectivity and the loss of information if they choose to only focus on the dominant peaks. SPA-STOCSY can set these separation boundaries with less ambiguity, by applying the intrinsic features of concentration heterogeneity among metabolites within the mixture. These features can help SPA-STOCSY separate small peaks from the dominant group if they are not from the same metabolites and offer a detailed annotation for the high-overlapping region.

However, SPA-STOCSY has some potential limitations and we have included some extra steps to address these intrinsic concerns. Firstly, SPA-STOCSY depends on the maximum homogeneity of the investigated samples (Alves *et al*., 2009). This suggests that the samples selected to explore structural connectivity should have the same metabolic changes and the peak locations should be well aligned. Specifically, we assume that the correlation pattern of the dataset directly reflects the strength of proton connectivity. If the spectra in the dataset are not well aligned or the phase is not properly corrected, the correlation pattern may not completely represent the structural information of molecules in the sample. To address this concern in our analysis of real samples, we used a chemical reference, DSS, a complex compound having a singlet at 0.00 ppm. The distances between peaks are relatively stable in all the samples, so when each sample is aligned to this singlet at 0.00 ppm, all spectra are homogeneous after the alignment. Although there is a chance that small positional variation may exist in the spectra, the SPA grouping mechanism can overcome it by a proper determination of window size *k* for the calculation of correlation landscape. As we use average correlation instead of point-to-point correlation, a tolerance of minor perturbation is embedded. Secondly, SPA captures the variation of signals and differentiates them from noise based on the assumption that the true concentrations of underlying metabolites are not identical across samples. This concern is easily solved with the intrinsic heterogeneity in real data, as there is no pair of biological samples that is identical.

Overall, SPA-STOCSY performs comparably to the widely used, operator-based Chenomx analysis in identifying candidate metabolites, but it eliminates the possibility of operator bias while returning the results in only seven minutes of computation time. It also outperforms SRV by capturing a higher percentage of the signal regions and the close-to-zero noise regions. Furthermore, summarizing the results from NMR spectra directly, SPA-STOCSY takes the library as a reference but is not limited only to the specific library. Unlike the previous identification methods (Hao *et al*., 2012; Ravanbakhsh *et al*., 2015; Lefort *et al*., 2019), SPA-STOCSY can offer the non-annotated metabolites by highlighting the correlated functional units with no match to any annotated metabolite. Accordingly, SPA-STOCSY brings new insights for metabolite compositions in diversified systems and offers researchers and clinicians a fast, highly accurate, and unbiased tool for NMR data analysis, with several key advantages over existing methods.

## Data availability

The data synthesized and generated for this study is available at https://github.com/LiuzLab/SPA_STOCSY

## Code availability

SPA-STOCSY is implemented with R. The codes and tutorials for SPA-STOCSY are available at https://github.com/LiuzLab/SPA_STOCSY

## Acknowledgement

We thank Cemal Karakas, MD, Soumya Patti, PhD, Marina Vannucci, PhD, and Christine Peterson, PhD for their helpful comments and inputs for this work; Advanced Technology Core support of NMR and Drug Metabolism core, the National Institute of General Medical Sciences (5R01GM120033 to M.M.S.). the Eunice Kennedy Shriver National Institute of Child Health & Human Development of the National Institutes of Health under Award Number P50HD103555 (M.M.S) for support of the Bioinformatics Core and the NMR and Drug Metabolism Core. The content is solely the responsibility of the authors and dose not necessarily represent the official views of the National Institutes of Health. The work was partially supported by the Cynthia and Antony Petrello Endowment (M.M.S.), the Chao Endowment (Z.L.) and the Huffington Foundation (Z.L.).

## Author contributions

M.M.S., Z.L., X.H. planned and initiated the project. M.M.S., Z.L. supervised the project. I.A-R and J.B. provided *Drosophila* tissue for analysis. L.M. performed the NMR experiments. X.H., W.W. designed and modified the algorithm, analyzed the data. G.A. provided guidance on statistical analysis. K.M. and D.W.Y. provided guidance on the NMR data collection. X.H., W.W., M.M.S wrote the manuscript.

## Corresponding authors

Correspondence to Mirjana Maletic-Savatic, Zhandong Liu.

