## Supplementary Figures for "SPA-STOCSY: An Automated Tool for Identification of Annotated and Non-Annotated Metabolites in High-Throughput NMR Spectra"

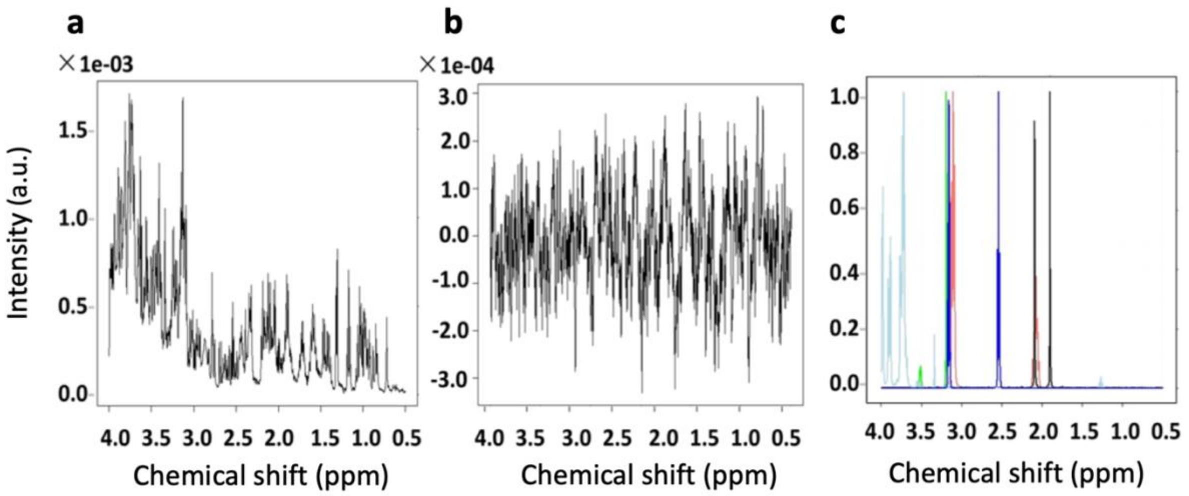


**Supplementary Fig. 1 | NMR simulation model. a**, A simulated spectrum with 50 metabolites. **b**, Simulated noise spectrum. **c**, A reference spectrum generated from five pure compound spectra downloaded from the BMRB database. The reference library data were Fourier Transformed, truncated to 0.5 to 4ppm region, and scaled to their highest peak. The intensity of the peaks is directly proportional to the metabolite concentration. Scaling enables manipulation of the pseudo concentration of a given metabolite in the simulation model.

| Scenarios \ Parameters | $\gamma$ | $\phi$ | $n$ | SNR_10 | SNR_30 | SNR_50 |
|---|---|---|---|---|---|---|
| 1 | 60 | 12 | 100 | 26.4 | 79.2 | 132 |
| 2 | 60 | 12 | 50 | 52.8 | 158.4 | 264 |
| 3 | 60 | 25 | 100 | 74.5 | 223.5 | 372.5 |
| 4 | 60 | 25 | 50 | 149 | 447 | 745 |

**Supplementary Table 1 |** Three sets of simulated datasets were generated containing 10, 30, or 50 metabolites. In each set, four scenarios were designed with different simulation parameters. (***Gamma, γ***) denotes the average concentration of metabolites in the sample; (***Phi, φ***) denotes the variance of different metabolite concentrations in the sample; ***n*** denotes the sample size. The SNR is calculated as the ratio of the variance of signals and the variance of the noise. With different simulation parameters, the SNR varies between each scenario and each set.

**Supplementary Table 2 | Automatic identification of metabolites from SPA-STOCSY for *Drosophila* data (detection threshold as 0.55) compared to Chenomx profiling results.**

In the separate PDF file.

**Supplementary Table 3 | Chenomx profiling results of 24 metabolites on hESCs spectra and their detection ratios in SPA-STOCSY**

| Metabolites identified with Chenomx in human cells | Total clusters | Number of clusters (0ppm to 4ppm) | Number of clusters identified | Detection ratio |
|---|---|---|---|---|
| 4-Aminobutyrate | 3 | 3 | 3 | 1 |
| Myo-inositol | 4 | 3 | 3 | 1 |
| Choline | 3 | 2 | 2 | 1 |
| O-Phosphocholine | 3 | 2 | 2 | 1 |
| Glycine | 1 | 1 | 1 | 1 |
| Lactate | 2 | 1 | 1 | 1 |
| Nicotinurate | 6 | 1 | 1 | 1 |
| Pyruvate | 1 | 1 | 1 | 1 |
| Leucine | 6 | 6 | 5 | 0.833 |
| Tyrosine | 5 | 3 | 2 | 0.667 |
| Proline | 7 | 6 | 3 | 0.5 |
| UDP-galactose | 15 | 4 | 2 | 0.5 |
| Valine | 4 | 4 | 2 | 0.5 |
| Alanine | 2 | 2 | 1 | 0.5 |
| Glutamate | 5 | 5 | 2 | 0.4 |
| Glutamine | 7 | 5 | 2 | 0.4 |
| Isoleucine | 6 | 6 | 2 | 0.333 |
| Phenylalanine | 6 | 3 | 1 | 0.333 |
| sn-Glycero-3-phosphocholine | 8 | 7 | 2 | 0.286 |
| DSS-d6 (Chemical Shape Indicator) | 1 | 0 | 0 | no peak in 0ppm - 4 ppm |
| Glucose-6-phosphate | 10 | 8 | 0 | not in library |
| AMP | 8 | 2 | 0 | not in library |
| UMP | 5 | 1 | 0 | not in library |
| Fumarate | 1 | 0 | 0 | not in library |
