## Supplementary Table 2 for "SPA-STOCSY: An Automated Tool for Identification of Annotated and Non-Annotated Metabolites in High-Throughput NMR Spectra"

| Candidate metabolites | SPA-STOCSY identified | Number of peaks ( 0 ppm to 4 ppm) | Number of peaks ( 4 ppm to 10 ppm) | Chenomx profiling | Comments |
|---|---|---|---|---|---|
| 1,3-Dihydroxyacetone | Y | 1 | 1 | Y | |
| 1,3-Dimethylurate | Y | 2 | 0 | N | Overlapping |
| 1,6-Anhydro-beta-D-Glucose | Y | 4 | 3 | Y | |
| 2-Ethylacrylate | N | 2 | 2 | Y | |
| 2-Hydroxyphenylacetate | Y | 1 | 4 | N | 1 peak between 0-4 (singlet) |
| 2-Hydroxyvalerate | Y | 5 | 1 | N | Not fit |
| 2-Methylglutarate | Y | 5 | 0 | N | Not fit |
| 2-Octenoate | Y | 5 | 2 | N | Overlapping |
| 2-Oxocaproate | Y | 3 | 0 | N | Not fit |
| 2-Oxoglutarate | Y | 2 | 0 | Y | |
| 2-Phosphoglycerate | Y | 2 | 1 | Y | |
| 3-Chlorotyrosine | Y | 3 | 3 | N | Overlapping |
| 3-Hydromuconate | Y | 1 | 1 | N | 1 peak between 0-4 (doublet) |
| 3-Hydroxy-3-methylglutarate | Y | 3 | 0 | N | Overlapping |
| 3-Hydroxybutyrate | Y | 3 | 1 | N | Not fit |
| 3-Hydroxyisovalerate | Y | 2 | 0 | N | Overlapping |
| 3-Hydroxykynurenine | Y | 1 | 4 | N | 1 peak between 0-4 (doublet) |
| 3-Hydroxyphenylacetate | Y | 1 | 4 | N | 1 peak between 0-4 (singlet) |
| 3-Methylxanthine | Y | 1 | 1 | N | 1 peak between 0-4 (singlet) |
| 4-Aminobutyrate | Y | 3 | 0 | Y | |
| 4-Aminohippurate | Y | 1 | 3 | N | 1 peak between 0-4 (doublet) |
| 4-Hydroxy-3-methoxymandelate | Y | 1 | 4 | Y | |
| 4-Hydroxyphenylacetate | Y | 1 | 2 | N | 1 peak between 0-4 (singlet) |
| 4-Pyridoxate | Y | 1 | 2 | N | 1 peak between 0-4 (singlet) |
| 5-Aminolevulinate | Y | 4 | 1 | N | Not fit |
| 5-Hydroxyindole-3-acetate | Y | 1 | 4 | N | 1 peak between 0-4 (singlet) |
| 5-Hydroxytryptophan | Y | 2 | 6 | N | Overlapping |
| 5-Methoxysalicylate | Y | 1 | 3 | N | 1 peak between 0-4 (singlet) |
| Acetamide | Y | 1 | 2 | Y | |
| Acetaminophen | Y | 1 | 2 | N | 1 peak between 0-4 (singlet) |
| Acetate | Y | 1 | 0 | Y | |
| Acetoacetate | N | 2 | 0 | Y | |
| Acetone | Y | 1 | 0 | Y | |
| Acetylsalicylate | Y | 1 | 4 | N | 1 peak between 0-4 (singlet) |
| Adenine | N | 0 | 2 | Y | |
| Adenosine | Y | 2 | 6 | Y | |
| Alanine | Y | 2 | 0 | Y | |
| AMP | N | 0 | 8 | Y | |
| Anserine | Y | 7 | 4 | N | Not fit |
| Arabinitol | Y | 7 | 0 | Y | |
| Arabinose | Y | 15 | 9 | N | Overlapping |
| Arginine | Y | 7 | 2 | Y | |
| Ascorbate | N | 2 | 2 | Y | |
| Asparagine | Y | 3 | 2 | N | Not fit |
| Aspartate | Y | 3 | 0 | Y | |
| beta-Alanine | N | 2 | 0 | Y | |
| Butyrate | Y | 3 | 0 | N | Not fit |
| Cadaverine | Y | 3 | 0 | N | Overlapping |
| Caffeine | Y | 3 | 1 | N | Overlapping |
| Caprate | Y | 9 | 0 | N | Overlapping |

| | | | | | |
|---|---|---|---|---|---|
| Caprylate | Y | 7 | 0 | N | Overlapping |
| Carnitine | Y | 5 | 1 | N | Overlapping |
| Carnosine | Y | 6 | 4 | N | Overlapping |
| Choline | Y | 2 | 1 | Y | |
| Cis-aconitate | Y | 1 | 1 | Y | |
| Citrate | N | 2 | 0 | Y | |
| Citrulline | Y | 7 | 1 | N | Not fit |
| Creatinine | Y | 1 | 1 | Y | |
| Cysteine | Y | 3 | 0 | Y | |
| Dimethyl sulfone | Y | 1 | 0 | Y | |
| Dimethylamine | Y | 1 | 0 | Y | |
| DSS | Y | 4 | 0 | Y | |
| dTTP | Y | 3 | 6 | N | Not fit |
| Ethanol | N | 2 | 0 | Y | |
| Ethanolamine | N | 2 | 0 | Y | |
| Ethylene glycol | Y | 1 | 0 | Y | |
| Ferulate | Y | 1 | 5 | N | 1 peak between 0-4 (singlet) |
| Fructose | Y | 11 | 3 | N | Overlapping |
| Fumarate | N | 0 | 1 | Y | |
| Galactarate | Y | 1 | 1 | Y | |
| Galactitol | Y | 3 | 0 | Y | |
| Galactonate | N | 5 | 1 | Y | |
| Glucarate | Y | 1 | 3 | N | 1 peak between 0-4 (triplet) |
| Glucitol | Y | 8 | 0 | Y | |
| Gluconate | Y | 4 | 2 | N | Overlapping |
| Glucose | Y | 12 | 2 | Y | |
| Glutamate | Y | 5 | 0 | Y | |
| Glutamine | N | 5 | 2 | Y | |
| Glutaric acid monomethyl ester | Y | 4 | 0 | N | Not fit |
| Glycerate | N | 2 | 1 | Y | |
| Glycerol | Y | 3 | 0 | Y | |
| Glycine | Y | 1 | 0 | Y | |
| Glycolate | Y | 1 | 0 | Y | |
| Guanidoacetate | Y | 1 | 0 | Y | |
| Guanosine | Y | 2 | 6 | N | Not fit |
| Hippurate | Y | 1 | 6 | N | 1 peak between 0-4 (doublet) |
| Histamine | Y | 2 | 2 | Y | |
| Histidine | Y | 3 | 2 | Y | |
| Homogentisate | Y | 1 | 2 | N | 1 peak between 0-4 (singlet) |
| Homoserine | Y | 2 | 0 | Y | |
| Homovanillate | Y | 2 | 3 | N | Overlapping |
| Indole-3-acetate | Y | 1 | 6 | N | 1 peak between 0-4 (singlet) |
| Inosine | Y | 2 | 6 | Y | |
| Isocitrate | Y | 3 | 1 | Y | |
| Isoleucine | N | 6 | 0 | Y | |
| Isopropanol | Y | 1 | 1 | N | Not fit |
| Isovalerate | Y | 3 | 0 | N | Not fit |
| Kynurenine | Y | 1 | 5 | N | 1 peak between 0-4 (doublet) |
| lactate | N | 1 | 1 | Y | |
| Lactose | Y | 24 | 4 | N | Overlapping |
| Leucine | N | 6 | 0 | Y | |
| Lysine | N | 7 | 0 | Y | |
| Malate | N | 2 | 1 | Y | |
| Malonate | Y | 1 | 0 | Y | |

| | | | | | |
|---|---|---|---|---|---|
| Maltose | Y | 24 | 4 | Y | |
| Mannitol | Y | 4 | 0 | Y | |
| Mannose | Y | 12 | 2 | N | Overlapping |
| Methanol | Y | 1 | 0 | Y | |
| Methionine | N | 5 | 0 | Y | |
| Methylamine | Y | 1 | 0 | N | 1 peak between 0-4 (singlet) |
| Methylguanidine | Y | 1 | 2 | N | 1 peak between 0-4 (singlet) |
| Myo-inositol | Y | 3 | 1 | N | Overlapping |
| N-Acetylaspartate | N | 3 | 2 | Y | |
| N-Acetylcysteine | Y | 3 | 2 | Y | |
| N-Acetylglutamate | Y | 4 | 2 | N | Not fit |
| N-Acetylglutamine | Y | 5 | 4 | Y | |
| N-Acetylserotonin | Y | 3 | 6 | N | Overlapping |
| N-Acetyltyrosine | Y | 3 | 4 | Y | |
| NADH | Y | 2 | 17 | N | Overlapping |
| Nicotinurate | Y | 1 | 5 | N | 1 peak between 0-4 (singlet) |
| N-Methylhydantoin | Y | 1 | 1 | Y | |
| O-Phosphocholine | Y | 2 | 1 | Y | |
| O-Phosphoethanolamine | N | 2 | 0 | Y | |
| O-Phosphoserine | Y | 1 | 3 | N | 1 peak between 0-4 (quadruplets) |
| Ornithine | Y | 5 | 0 | N | Not fit |
| Pantothenate | Y | 8 | 1 | N | Not fit |
| p-Cresol | Y | 1 | 2 | N | 1 peak between 0-4 (singlet) |
| Phenylacetate | Y | 1 | 3 | N | 1 peak between 0-4 (singlet) |
| Phenylalanine | Y | 3 | 3 | N | Overlapping |
| Proline | N | 6 | 1 | Y | |
| Pyridoxine | Y | 1 | 3 | Y | |
| Pyruvate | Y | 1 | 0 | Y | |
| Ribose | Y | 15 | 9 | N | Overlapping |
| Salicylurate | Y | 1 | 5 | N | 1 peak between 0-4 (singlet) |
| Sarcosine | N | 2 | 0 | Y | |
| Serine | Y | 3 | 0 | Y | |
| Serotonin | Y | 2 | 5 | N | Overlapping |
| sn-Glycero-3-phosphocholine | Y | 7 | 1 | Y | |
| S-Sulfocysteine | N | 2 | 1 | Y | |
| succinate | Y | 1 | 0 | Y | |
| Sucrose | Y | 11 | 3 | N | Overlapping |
| Taurine | Y | 2 | 0 | Y | |
| Threonate | Y | 3 | 1 | Y | |
| Threonine | N | 2 | 1 | Y | |
| Theophylline | Y | 2 | 1 | N | Overlapping |
| Thymine | Y | 1 | 1 | N | 1 peak between 0-4 (singlet) |
| Thymol | Y | 3 | 3 | Y | |
| trans-4-Hydroxy-L-proline | Y | 4 | 2 | N | Overlapping |
| trans-Aconitate | Y | 1 | 1 | N | 1 peak between 0-4 (singlet) |
| Trimethylamine | Y | 1 | 0 | Y | |
| Trimethylamine N-oxide | Y | 1 | 0 | Y | |
| Tropate | Y | 2 | 4 | N | Overlapping |
| Tryptophan | Y | 2 | 7 | N | Overlapping |
| Tyrosine | Y | 3 | 2 | N | Overlapping |
| UDP-glucuronate | Y | 3 | 10 | N | Not fit |
| Valine | N | 4 | 0 | Y | |
| Vanillate | Y | 1 | 3 | N | 1 peak between 0-4 (singlet) |
| Xylose | Y | 10 | 2 | N | Overlapping |
